# Domestication effects on immune response: comparison of cell-mediated immune competence in wild and domesticated Bengalese finch

**DOI:** 10.1101/2021.02.24.432813

**Authors:** Kenta Suzuki, Kazuo Okanoya

## Abstract

Domesticated Bengalese finch (*Lonchura striata* var. *domestica)* lack natural selection pressures and consequently have more complex songs and altered behavioural and psychological traits when compared to their wild ancestor, the white-rumpud munia (*Lonchura striata*). Clarifying the sexual traits and life history trade-offs in Bengal finches will be help to improve our understanding of the evolution of complex songs and domesticated traits. Here, we have assessed the immune competence of the Bengalese finch and the white-rumped munia using phytohemagglutinin (PHA) tests to create an index of cell-mediated immune responses. We found that the Bengalese finch had a significantly larger immunocompetence than the white-rumped munia, indicating that they devote more resources to both immunity and reproduction. Thus, there is no trade-off but a positive relationship between immunocompetence and reproductive traits, which may be related to the release from natural selection pressures. These results will be useful in understanding the mechanisms by which domestication-induced changes due to a lack of natural selection pressure affect behavioural and physiological changes.

## 1. Introduction

The Bengalese finch (*Lonchura striata* var. *domestica*) is a domesticated strain of the wild strains, white-rumped munia (*Lonchura striata*) (Washio, 1996; Okanoya 2004a, b). It evolved phonologically and syntactically complex songs, which are considered sexual traits (Honda and Okanoya, 1999; Okanoya 2004a, b). Domestication processes have also altered the physiological and behavioural traits of the Bengalese finch. The white-rumped munia have traits that are advantageous for survival in the wild, such as higher fear reactivity, aggression, and corticosterone levels, when compared with the Bengalese finch (Suzuki et al., 2012; 2013; 2014a, b, and unpublished data). It is likely that the latter has developed reproductive traits, rather than the traits that are advantageous in the wild due to the release from natural selection pressures and the presence of artificial selection. Incidentally, the song traits of the Bengalese finch have never been artificially selected by humans (Washio, 1996).

There are several possible hypotheses for the relationship between the evolution of sexual traits and life history trade-offs in the Bengalese finch. First, in the wild environment, high immunocompetence must be maintained to protect against the high risk of infection but in the domesticated strains, there is no risk in the rearing environment, so the cost to immunity may be reduced and sexual traits can be developed. Selected lines of domestic fowl (*Gallus gallus domesticus*) with different immunoresponsiveness levels showed that a high response line had a smaller comb size as a sexual ornament (Verhulst et al., 1999). Thus, there appears to be a negative correlation between immunocompetence and sexual traits. Second, the immunocompetence handicap hypothesis (ICHH) should be considered (Folstad & Karter, 1992). It is known that testosterone develops sexual traits and suppresses immunocompetence. Males that have developed sexual traits and survive without pathogen infections are superior with high immunocompetence. Therefore, sexual traits are considered an indicator of high immunocompetence. For example, there is evidence that male house finches (*Carpodacus mexicanus*) that survived an epidemic had significantly redder plumage, a sexual trait, than the males that did not survive (Nolan et al., 1998). Third, the developmental stress hypothesis suggests that song complexity is related to stress conditions during development (Buchanan et al., 2003), and since stress lowers immunocompetence and sexual traits, males with lower stress levels may be able to maintain higher immunocompetence and develop complex song traits as attractive sexual traits.

By comparing the immunity of Bengalese finch and white-rumped munia, it is thought that the evolutionary mechanisms of the domesticated traits (i.e., complex song evolution) may be better understood. Therefore, the present study aimed to compare the immune competence between domesticated Bengalese finches and their wild ancestor, the white-rumped munia. Phytohemagglutinin (PHA) tests were performed to index the cell-mediated immune responses. The PHA test is practical and reliably measures immunocompetence as acquired T-cell mediated immunocompetence (Tella et al., 2008).

## 2. Method

### 2. 1. Subjects and husbandry

Thirty-six adult Bengalese finches (*Lonchura striata* var. *domestica*, 19 males and 17 females) and 38 adult white-rumped munia (*Lonchura striata*, 18 males and 20 females) were used for the experiment. The Bengalese finches were bred in our laboratory. White-rumped munias were bought from a commercial supplier (n = 12), captured in the wild in Taiwan (n = 3), or bred in our laboratory (n = 23). The birds were housed in groups of four (each species and sex) in the same type of stainless-steel cages (cage size: 370 mm × 415 mm × 440 mm, equipped with two wooden perches) in a common room at RIKEN Brain Science Institute (BSI) with a 13-/11-h light/dark cycle; the birds were supplied food and water. The ambient room temperature was approximately 25 °C with 50% humidity. Seed mixture, shell grit, and vitamin-enhanced water were available *ad libitum*. All experimental procedures and the housing conditions of the birds were approved by the Animal Experiments Committee at RIKEN (#H24-2-229), and all of the birds were cared for in accordance with the Institutional Guidelines for Experiments Using Animals.

### 2. 2. PHA test

Phytohemagglutinin (PHA) responses (wing-web swelling test) are commonly used to assess cell-mediated immune competence in passerine birds, as it is a T-cell stimulant (Martin et al., 2006; Tella et al., 2008). Before the injection, we cleaned the left-wing web of the birds with an alcohol swab, and measured its thickness four times using a digital spessimeter (Mitsutoyo, Kanagawa, Japan) to the nearest 0.001 mm. Then, 0.03 mL of 5 mg/mL solution of PHA-P (Cat. L8754, Sigma Chemical Co., St Louis, MO, USA) was dissolved in phosphate-buffered saline (1 × PBS) and injected into the left-wing-web (patagium) using a 27-gauge needle. Four measurements of the wing web thickness at the injected point were taken after 24 h. The mean of these measured values was used for the analysis. The PHA index (PHA response) was calculated as the difference between the post injection and the pre-injection values for the wing-web thickness.

### 2. 3. Statistical analysis

The PHA indexes of the Bengalese finch and white-rumped munias were analysed using two-way analysis of variance (ANOVA, factors: strain and sex). Comparison of the PHA indices in the three groups of white-rumped munias, under different rearing conditions (bred, bought, and captured individuals), were also assessed using ANOVA. Statistical analyses were performed using Stat View software (version 5, SAS Institute Inc., Cary, NC, USA). Values of *p* < 0.05 were considered significant.

## 3. Results

The PHA index (Fig. 1) showed that there were significant main effects for the strain [*F* (1, 70) = 37.52, *p* < 0.0001], but no significant main effects for sex [*F* (1, 70) = 1.398 × 10^−6^, *p* = 0.999] or interactions between strain and sex [*F* (1, 70) = 3.733, *p* = 0.06]. The PHA index was significantly higher in the Bengalese finches [mean = 0.846, standard error of mean (SEM) = 0.039] than in the white-rumped munias (mean = 0.557, SEM = 0.029). We found no significant differences in the PHA indexes between the different rearing conditions (bred, bought, and captured individuals) for the white-rumped munias [*F* (2, 35) = 0.188, *p* = 0.830].

**Fig. 1.**
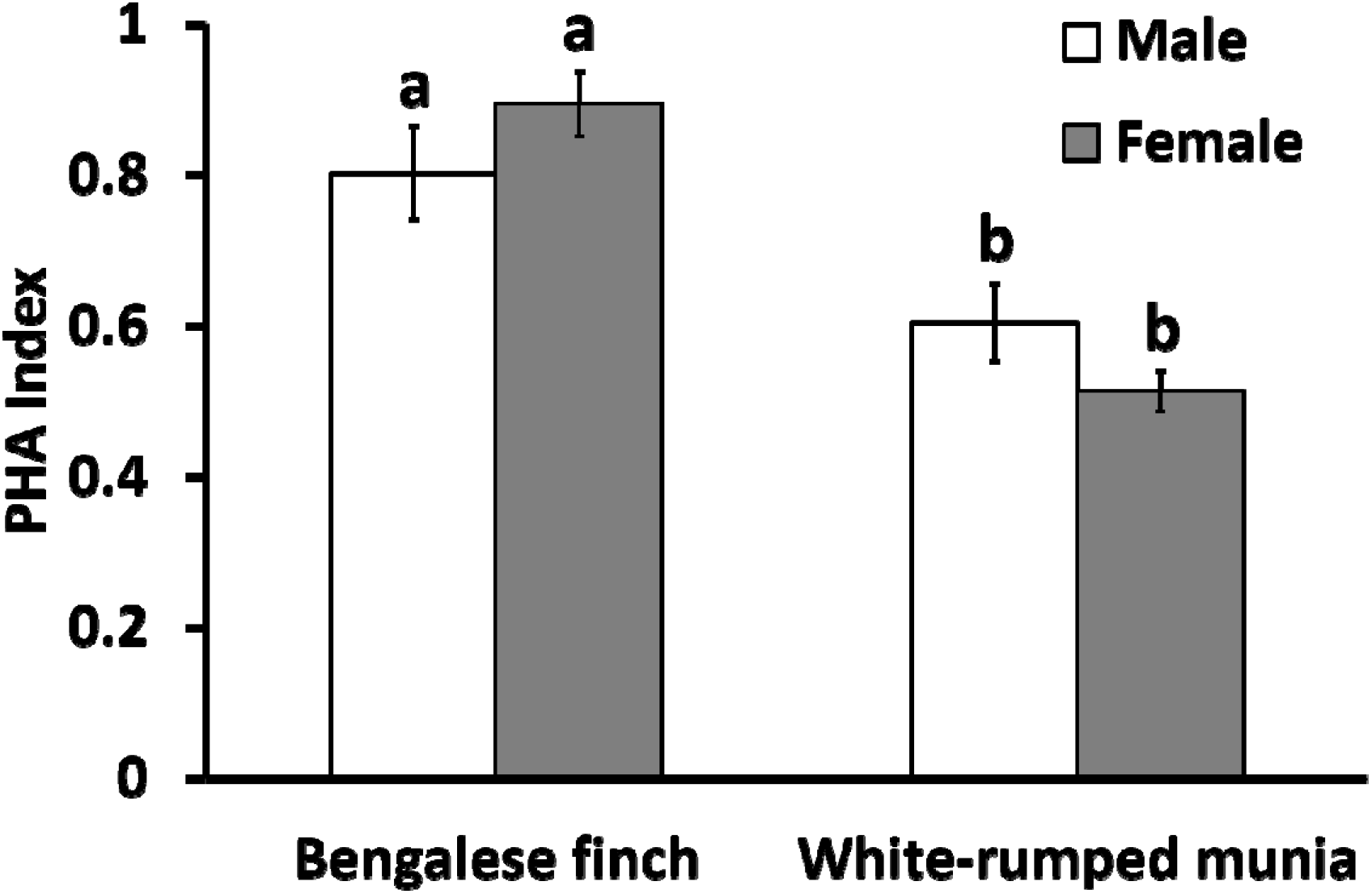
Phytohemagglutinin (PHA) index in Bengalese finches (n = 36) and white-rumped munias (n = 38). Bars are means, and vertical lines are standard error of mean (SEM). Open bars represent male birds, and closed bars represent female birds. Different letters indicate significant differences. The PHA indexes of the Bengalese finch and white-rumped munias were analysed using two-way analysis of variance (ANOVA, factors: strain and sex). Differences were found in the PHA index between Bengalese finches and white-rumped munias (*P* < 0.0001). No sex differences (*P* > 0.05) were observed in the PHA indexes.

## 4. Discussion

The PHA test was used to compare the immune capacities of the domesticated Bengalese finch and its wild ancestor, the white-rumped munia, and found that the former had a significantly larger immunocompetence than the latter. The results showed that the immune capacity and sexual traits were both higher in the domesticated Bengalese finch than the wild strain. An investigation of zebra finches (*Taeniopygia guttata*), also found that the domestic birds had a stronger PHA response than the wild birds (Tschirren et al., 2009). Furthermore, the domestication process also seemed to improve the immune status of fish, as F4 tamed Eurasian perch juveniles displayed higher immune capacities than the F1 (*Perca fluviatilis*; Douxfils et al., 2011). These results indicate that domestication enhances immunity in a variety of species. The laboratory-bred Bengal finches and white-rumped munia were raised under the same rearing conditions and had similar durations and levels of human contact. The results of this study did not show any differences in the PHA responses among the different rearing conditions (bred, bought, and captured individuals) used for the white-rumped munias. Therefore, it is suggested that the PHA of the Bengalese finches was higher due to domestication, irrespective of the differences in their rearing conditions.

The domesticated Bengalese finches exhibit a high-level immune response and complex song traits compared with the wild birds. Therefore, there does not appear to be a trade-off between immune capacity and sexual traits. Testosterone is known to develop sexual characteristics and suppress the immune system. According to the immunocompetence handicap hypothesis (ICHH), sexual traits are considered an indicator of high immunocompetence because males that have developed sexual traits and survive without pathogen infections are superior with high immunocompetence (Folstad & Karter, 1992). It is possible that the Bengalese finch could maintain a highly competent immune system even with the use of high testosterone level in the development of its complex song trait as there is no pressure of natural selection. However, there is preliminary data showing that there is no difference in the testosterone levels between Bengal finches and white-rumped munias (Tobari et al., 2019). Therefore, the immunocompetence handicap hypothesis (ICHH) does not seem to apply. Finally, the stress level (faecal corticosterone concentration) during the developmental periods was lower in the Bengalese finch than in the white-rumped munias (Suzuki et al., 2014a, b), and this resulted in improved development of immunocompetence and sexual traits. Previous studies into the repertoire size and song bout durations of European starlings (*Sturnus vulgaris*) showed that developmental stress decreases song complexity, and there was also a positive relationship with song traits and immune response (PHA response; Buchanan et al., 2003; Spencer et al., 2004). This study suggests that the results are consistent with the developmental stress hypothesis (Buchanan et al., 2003).

The mechanisms by which domestication alters the immune responses are not clear. Differences in allele frequencies for the major histocompatibility complex (MHC), whose molecules play a crucial role in adaptive immune responses, have been identified between wild zebra finch subspecies and domesticated populations, which may indicate immune adaptations to pathogen pressures (Newhouse and Balakrishnan, 2015). However, further research is required to determine whether these changes actually affect the differences in immune capacity.

It is possible that corticosterone will affect immunocompetence. Faecal corticosterone levels were previously found to be lower in Bengal finches than in wild strains (Suzuki et al., 2012), despite their higher PHA responses. Butler et al. (2010) reported that there was a negative relationship between corticosterone and PHA responses in sham-implanted American kestrel (*Falco sparverius*) nestlings. Birds that naturally maintain higher levels of circulating corticosterone may also have reduced levels of cutaneous immune function (Butler et al., 2010). Moreover, house sparrow (*Passer domesticus*) nestlings injected with corticosterone showed a weaker immune response (PHA response) than the controls (Loiseau et al., 2008). There are also reports that corticosterone has an immunosuppressive effect (Buchanan 2000). Bengalese finch may maintain high immune capacities as corticosterone levels decrease due to domestication.

Zebra finch studies that used a selection line with peak corticosterone levels for the stress responses, found that peak corticosterone titers had a positive effect on the PHA response. The selection line with high peak corticosterone levels had a higher PHA response (Roberts et al., 2007). In the American kestrel study, corticosterone-treated nestlings showed an increased amount of swelling (immunoresponsiveness) after the increase in corticosterone exposure had ended, when compared with that in the controls (Butler et al., 2010). The seemingly different effects of corticosterone on immunity (immunostimulatory or immunosuppressive) may be due to the differences in low-level or high-level and acute or chronic responses, and differences in the effects of the two types of corticosterone receptors: mineralocorticoid and glucocorticoid. Low-levels of corticosterone may enhance immunocompetence after hormone exposure, while chronically elevated corticosterone has immunosuppressive effects (Butler et al., 2010).

In summary, these results indicate that domesticated Bengal finches have a higher response to PHA than their wild ancestors, the white-rumped munias do. This suggests that the domestication of Bengal finches allowed them to devote more resources to both immunity and reproduction. Thus, there is no trade-off but a positive relationship between immunity and reproductive traits, which may be related to the release from natural selection pressures and the decrease in stress hormones due to domestication. These results will be useful in understanding the mechanisms by which domestication-induced changes in selection pressure affect behavioural and physiological changes, leading to the formation of domestication traits (domestic syndromes). The details of these mechanisms however need to be further investigated in the future.

## Acknowledgements

We thank Dr. Yuko Ikkatai and Ms. Keiko Asai (RIKEN Brain Science Institute, Tokyo, Japan) for animal care. This study was supported by JST-ERATO and RIKEN Brain Science Institute.

## Funding

This research receives grants from JSPS KAKENHI (15K14581, 17H06380, 20H00105) to KO.

## Notes

### Competing Interest Statement

The authors have declared no competing interest.

